# A Closed-Loop Mathematical Structure of Mechanics–Turnover Coupling for Mechanical Adaptation in Living Systems

**DOI:** 10.64898/2026.02.20.707125

**Authors:** Eiji Matsumoto, Shinji Deguchi

## Abstract

Mechanical adaptation underlies mechanical homeostasis by allowing living systems to restore characteristic mechanical variables under sustained perturbations. Across biological scales, turnover-mediated remodeling enables mechanical adaptation by continuously renewing internal structures under load. Despite extensive progress in this field, it remains to be established what closed-loop mathematical structure of mechanics–turnover coupling is sufficient to guarantee homeostasis and how the characteristic adaptation timescale emerges from this coupling. Here, we identify the minimal mathematical structure of closed-loop mechanics–turnover coupling, providing a unifying description of mechanically adaptive remodeling across scales. We derive an analytical expression for the adaptation timescale as a function of the coupling between internal mechanical parameters and turnover kinetics, enabling direct cross-system comparison. To isolate this structure, we formulate a dynamical model linking mechanics and turnover, and establish conditions under which the closed-loop dynamics exhibit integral action. Specifically, our model describes how deviations in the mechanical state modulate the turnover of an internal structural state, and the renewed structure feeds back onto mechanics in a negative-feedback direction, driving recovery toward a reference state. We define systems satisfying this structure as *Feedback Adaptive Turnover-mediated Environment-Dependent (FATED)* systems. As an experimental example, we formulate mechanical adaptation in terms of mechanically regulated actin turnover. With the generalization of this architecture, we evaluate cross-system consistency by comparing reported adaptation and turnover timescales across representative remodeling systems.

## 1 Introduction

Homeostasis refers to the capacity of living systems to maintain functional stability under internal and external perturbations, a concept described by Cannon as the regulation of the internal state (Cannon 1939). Among the homeostatically regulated variables, mechanical quantities such as stress, strain, tension, pressure, and force are also maintained near characteristic levels. This principle is exemplified by Wolff’s law (Wolff 1986), which states that bone remodels under mechanical loading to restore an appropriate mechanical state. More generally, such phenomena are referred to as mechanical homeostasis. Mechanical adaptation denotes the stabilization of a regulated mechanical quantity through structural remodeling of load-bearing systems. Mechanical adaptation is essential for development, long-term tissue maintenance, and preservation of functional stability, in which mechanical factors regulate tissue architecture, morphogenesis, and cellular signaling (Mammoto et al. 2013; DuFort et al. 2011; Hahn and Schwartz 2009; Liu et al. 2022). When mechanical regulation operates properly, living systems adapt to altered loading and restore the regulated mechanical state. By contrast, disruptions of key regulatory molecules or signaling pathways can impair the feedback mechanisms that normally stabilize mechanical variables. Persistent mechanical dysregulation in turn has been linked to sustained pro-inflammatory signaling and chronic immune stimulation (Hahn and Schwartz 2009; Chien 2007; Humphrey and Schwartz 2021; Kaunas and Deguchi 2011). Together, these findings highlight mechanical adaptation as a broadly important process across scales.

In hierarchically organized living systems, these structures are dynamic and continually renewed through turnover. At molecular and cellular levels, cytoskeletal networks polymerize, depolymerize, and exchange subunits, allowing mechanically relevant architecture to be updated (Pollard and Cooper 2009; Fletcher and Mullins 2010; Saito et al. 2022; Buenaventura et al. 2024). At tissue scales, stress-dependent growth, constrained-mixture turnover, and cell turnover replace constituents (Rodriguez et al. 1994; Humphrey 2021; Ganusov et al. 2005; Sender and Milo 2021). In bone, cyclic resorption and formation renew material and contribute to maintaining mechanical integrity (Bolamperti et al. 2022). Across these scales, turnover therefore provides a generic physical mechanism for renewing internal structure under sustained load.

Despite extensive studies of remodeling laws and mechanistic pathways across scales, it remains unclear what essential mathematical structure of the closed-loop mechanics–turnover coupling is sufficient to guarantee mechanical homeostasis. Here, we seek an essential state-and-coupling description that is independent of system-specific constitutive details. Many studies have focused on either system-specific molecular mechanisms or constitutive formulations that map mechanical stimuli to deposition, degradation, growth, or other remodeling responses (Rodriguez et al. 1994; Humphrey 2021; Ambrosi and Guana 2007). Established growth-and-remodeling and constrained-mixture frameworks typically specify system-dependent remodeling laws. However, these descriptions do not make explicit the general closed-loop mathematical structure with negative feedback shared across scales (Fig. 1). In particular, turnover alone does not uniquely determine which mechanical quantities are maintained near a reference level. Even if remodeling nominally provides negative feedback, the system can still exhibit persistent offsets, drift, overshoot, or instability. Thus, existing descriptions have yet to identify the general structural conditions under which turnover-mediated renewal ensures adaptation. It also remains unclear how the characteristic adaptation timescale emerges from the coupling between internal mechanical parameters and turnover kinetics.

**Fig. 1.**
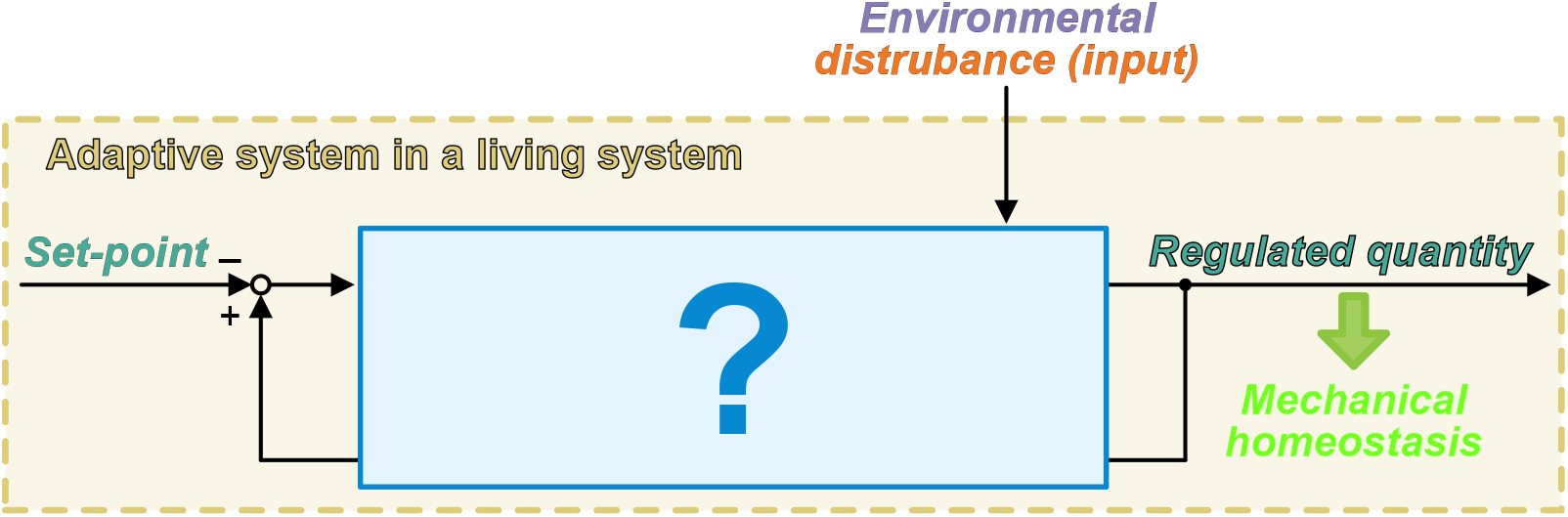
What is the essential closed-loop mathematical structure that enables turnover-mediated remodeling to achieve mechanical adaptation?

Here, we identify the minimal mathematical structure of closed-loop mechanics–turnover coupling that is necessary to ensure turnover-mediated homeostasis, thereby providing a unifying understanding of mechanically adaptive remodeling across scales. We then obtain an analytical expression for the adaptation timescale as a function of the coupling between internal mechanical parameters and turnover kinetics, enabling direct cross-system comparison through the resulting characteristic adaptation time constant. To isolate this closed-loop structure, we formulate a dynamical model linking mechanics and turnover, without invoking a control-theoretic approach. Under a local linearization, we show that the closed loop has integral action for homeostasis. This property explains why turnover-mediated remodeling can achieve adaptation to disturbances. We refer to closed-loop systems satisfying this structure as *Feedback Adaptive Turnover-mediated Environment-Dependent (FATED)* systems, characterized by structural renewal that closes the mechanics–turnover loop and shapes adaptation to changes in the mechanical environment.

As a simple experimentally grounded case, we first formulate a remodeling model of mechanically regulated actin turnover. We then generalize this class of closed-loop systems to describe *FATED* systems. Finally, we assess cross-system consistency by comparing reported adaptation and turnover timescales across representative remodeling systems.

## 2 Remodeling model for mechanically regulated actin turnover

### Force-dependent actin turnover in *in vitro* reconstitution experiments

*In vitro* reconstitution experiments have shown that mechanical loading modulates actin turnover and promotes filament elongation across diverse force-application modalities, including microfluidic drag, optical and magnetic tweezers, and myosin-based biomimetic assays (Zimmermann and Kovar 2019; Jégou et al. 2013). In the presence of profilin and formins, e.g., mDia1, tension enhances formin-mediated polymerization, resulting in sustained filament elongation under load (Courtemanche et al. 2013; Yu et al. 2017). From a mechanical viewpoint, loading increases the stress carried by the filament. Under fixed-end conditions, the end-to-end length is prescribed. In a one-dimensional elastic description, stress is determined by the mismatch between this imposed length and an internal stress-free rest length. Turnover-driven elongation therefore reduces the elastic extension relative to the rest length and relaxes stress even under persistent deformation. We represent this mechanically as an increase in an effective stress-free rest length and model turnover-mediated elongation as a rest-length update. This description makes explicit a closed mechanical loop in which elevated stress promotes rest-length elongation, which in turn reduces stress (Fig. 2). This feedback structure provides a minimal realization of mechanically regulated structural renewal under persistent load. The reconstituted actin system enables isolation of the adaptive loop without additional regulatory complexity.

**Fig. 2.**
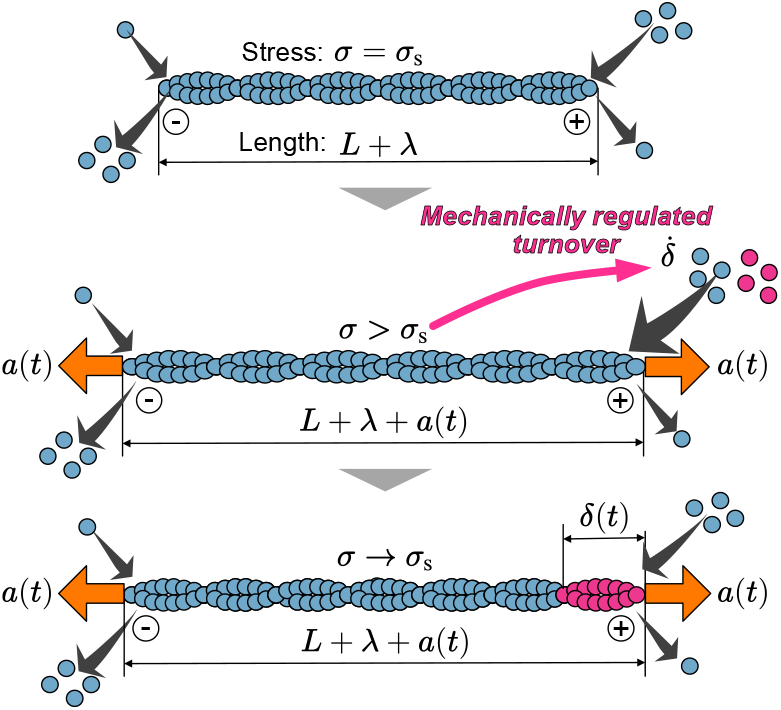
Problem setting for mechanically regulated actin adaptation. An imposed deformation *a*(*t*) under fixed-end boundaries perturbs a minimal single-filament actin model from a mechanically homeostatic reference state with *σ* = *σ*_s_ and baseline deformation *λ*. The resulting stress deviation, *σ > σ*_s_, shifts the filament turnover rate from its baseline level, producing a stress-dependent update of the rest length, *δ*(*t*). This rest-length change relaxes the stress back toward the set-point while the deformation *a*(*t*) persists.

### Elastic constitutive law

As a mechanical description, we model a single actin filament (AF) as a linear elastic element. We use a Hookean constitutive relation between the mechanical stress *σ*(*t*) and strain *ε*(*t*),

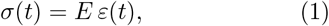

where *E* denotes the Young’s modulus of the AF.

### Kinematics

We formulate a minimal model of mechanically regulated actin adaptation under the fixed-end deformation setup (Fig. 2). At *t* = 0, the AF is in a mechanically homeostatic reference state with stress set-point *σ* = *σ*_s_ and baseline deformation *λ*. We denote the initial rest length by *L* and define the turnover-induced change in the rest length by *δ*(*t*), so that the rest length becomes *L*+*δ*(*t*). Upon an externally applied end-to-end deformation *a*(*t*), the end-to-end length is maintained at *L*+*λ*+*a*(*t*), while turnover updates the rest length to *L* + *δ*(*t*).

Hence, the elastic extension is *λ* + *a*(*t*) − *δ*(*t*), and the strain is

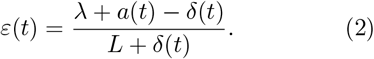

In the homeostatic reference state at *t* = 0, the strain and stress are

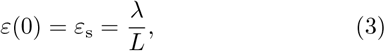

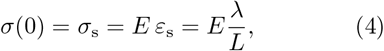

respectively. here, *ε*_s_ denotes the set-point of strain.

### Stress-regulated turnover as a rest-length update

We model mechanically regulated turnover as a stress-dependent change in the rest length. Stress deviations from the set-point shift the turnover rate from its baseline level, and thereby update the rest length, as observed in *in vitro* experiments. To capture the intrinsic behavior, we adopt a linear relationship

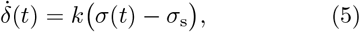

where *k* is the mechano-chemical coupling coefficient, relating stress deviation to the turnover-mediated rest-length update. We define *a*(*t*) and *δ*(*t*) as deviations from the homeostatic reference state, such that *a*(0) = 0 and *δ*(0) = 0. Since *σ*(0) = *σ*_s_ at homeostasis, Eq. (5) implies 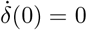.

### Linearization and derivation of the characteristic adaptation time constant

To extract the characteristic adaptation time, we linearize the model around the homeostatic reference state. The only nonlinearity appears in the kinematic relation (Eq. (2)), which we expand to first order around *δ*(*t*) = 0. Assuming *L*≫ *λ, a*(*t*), *δ*(*t*), we obtain

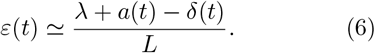

Using the elastic relation (Eq. (1)), the stress is expressed as

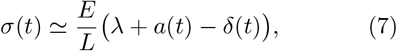

so that the stress error becomes

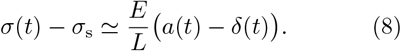

Substituting this expression into the turnover law (Eq. (5)), we obtain a linear equation governing the rest-length change in response to the imposed deformation,

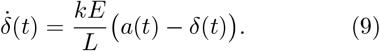

This equation describes first-order relaxation dynamics of *δ*(*t*) toward the imposed deformation *a*(*t*). The characteristic adaptation time constant is therefore given by

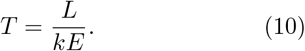

This time constant sets the rate of adaptation to an imposed deformation.

### Representative temporal responses

We show the step response of the nonlinear model as the simplest dynamical behavior, obtained by numerical integration of Eq. (1)–Eq. (5) (Fig. 3). This demonstrates complete adaptation in the sense that the stress error *σ*(*t*) *− σ*_s_ returns to zero under a sustained step deformation. For deformations within the linear regime, the adaptation time constant is defined by Eq. (10).

**Fig. 3.**
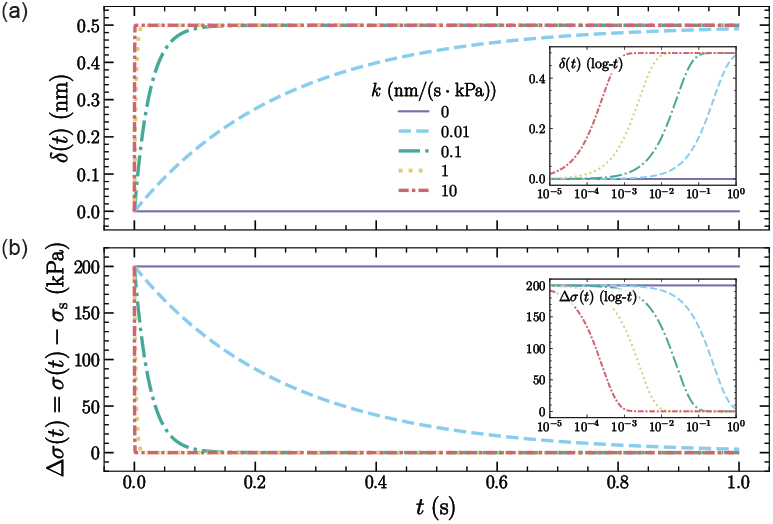
Step response of the minimal model. (a) The turnover-induced rest-length change *δ*(*t*) accelerates as the coupling coefficient *k* increases, whereas *k* = 0 yields no adaptive update. (b) The stress error Δ*σ*(*t*) = *σ*(*t*) *− σ*_s_ decays toward zero for *k >* 0. In the linearized model, the adaptation time constant is *T* = *L/*(*kE*); increasing *k* reduces *T* while preserving the steady-state value of the rest length and stress.

In the linearized model, the step response admits a closed-form solution. Under a step deformation *a*(*t*) = *a*_0_*H*(*t*), where *H*(*t*) is the Heaviside step function, *H*(*t*) = 0 for *t <* 0 and *H*(*t*) = 1 for *t* ≥ 0, the rest-length update is given by

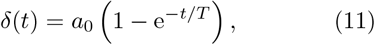

while the stress error decays exponentially as

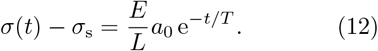

Varying *k* directly sets the time constant *T* while leaving the steady-state solution unchanged, as reflected by the different relaxation rates (Fig. 3). All curves were generated using the parameter values listed in Table 1. The step deformation amplitude *a*_0_ is estimated from reported force scales in formin-assisted actin polymerization experiments by converting an axial force perturbation into an effective elongation using the linear elastic relation *F* = *Ka*_0_, with *K* = *ES/L*. The ramp response is discussed in Appendix 1.

**Table 1.** Parameters used for Fig. 3. The model is driven by a step deformation of amplitude *a*_0_. Derived quantities are included for completeness.

| Quantity | Symbol | Value | Reference |
| --- | --- | --- | --- |
| Filament length | $L$ | 5 $\mu\text{m}$ | (Burlacu et al. 1992; Sept et al. 1999; Jégou et al. 2013) |
| Young’s modulus | $E$ | 2 GPa | (Gittes et al. 1993; Kojima et al. 1994; Isambert et al. 1995; Shamloo and Mehrafrooz 2018a) |
| Cross-sectional area | $S$ | 38.5 $\text{nm}^2$ | Derived from $S = \pi(d/2)^2$ with $d = 7 \text{ nm}$ (Alberts et al. 2022) |
| Step deformation amplitude | $a_0$ | 0.5 nm | Estimated from reported force scales (Courtemanche et al. 2013; Jégou et al. 2013; Kubota et al. 2017; Yu et al. 2017; Kubota et al. 2018; Yu et al. 2018)<br>by converting these force into a deformation amplitude using $a_0 = F_0/K$ with $K = ES/L$ |
| Mechano-chemical coupling coefficient | $k$ | 0, 0.01, 0.1, 1, 10 $\text{nm}/(\text{s} \cdot \text{kPa})$ | Explored parameter range from (Courtemanche et al. 2013; Jégou et al. 2013; Kubota et al. 2017; Yu et al. 2017; Kubota et al. 2018; Yu et al. 2018) |
| Relative deformation | $a_0/L$ | $1.0 \times 10^{-4}$ | Derived |
| Initial stress error | $\Delta\sigma(0^+) = Ea_0/L$ | 200 kPa | Derived |
| Adaptation time constant | $T = L/(kE)$ | $\infty, 0.25, 0.025, 0.0025, 0.00025 \text{ s}$ | Derived |

## 3 System interpretation and integral action

### Linearized dynamics and Laplace-domain representation

We interpret the linearized model in Sec. 2 within the framework of feedback control theory. Defining the stress error as Δ*σ*(*t*) := *σ*(*t*) *−σ*_s_, the turnover law (Eq. (5)) implies

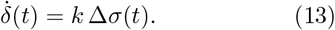

Let *a*(*s*), Δ*σ*(*s*), and *δ*(*s*) denote the Laplace transforms of *a*(*t*), Δ*σ*(*t*), and *δ*(*t*), respectively. Taking the Laplace transform of Eq. (13) with *δ*(0) = 0 yields

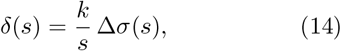

which corresponds to an integrator *k/s*. In the linearized kinematics (Eq. (6)), the strain is written in Laplace form as

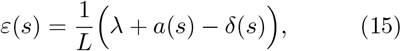

where *λ* is a constant offset that sets the homeostatic stress. The constitutive law is expressed as

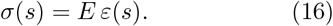

Since *σ*_s_ = *Eλ/L*, the offset *λ* cancels when forming Δ*σ*(*s*) = *σ*(*s*) *− σ*_s_. Using Eq. (15)–Eq. (16) together with Eq. (14), the closed-loop relation between the deformation input and the stress error can be written as

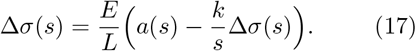

The stress response is described by the closed-loop transfer function

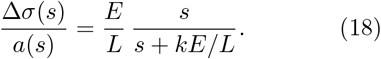

This transfer function makes explicit (i) the single relaxation timescale set by the pole at *s* = *−kE/L*, and (ii) complete adaptation to step deformations due to the zero at the origin. A formal proof of this property using the final value theorem is provided (Appendix 2).

### Block diagram interpretation

The block diagram (Fig. 4) provides a system-level interpretation of the actin remodeling model by making its feedback architecture explicit. The turnover law appears as an integrator *k/s* acting on the stress error Δ*σ*(*s*), and the resulting state update enters the mechanical relation with a negative sign, closing a negative-feedback loop. This combination of integral action and negative feedback is sufficient to ensure complete adaptation under local linearization and closed-loop stability. This interpretation isolates the architecture responsible for adaptation, independent of system-specific details, and clarifies why turnover-mediated state renewal yields integral action. It also provides a template for the next section, where we generalize it to define a closed-loop mathematical structure for mechanical adaptation across systems.

**Fig. 4.**
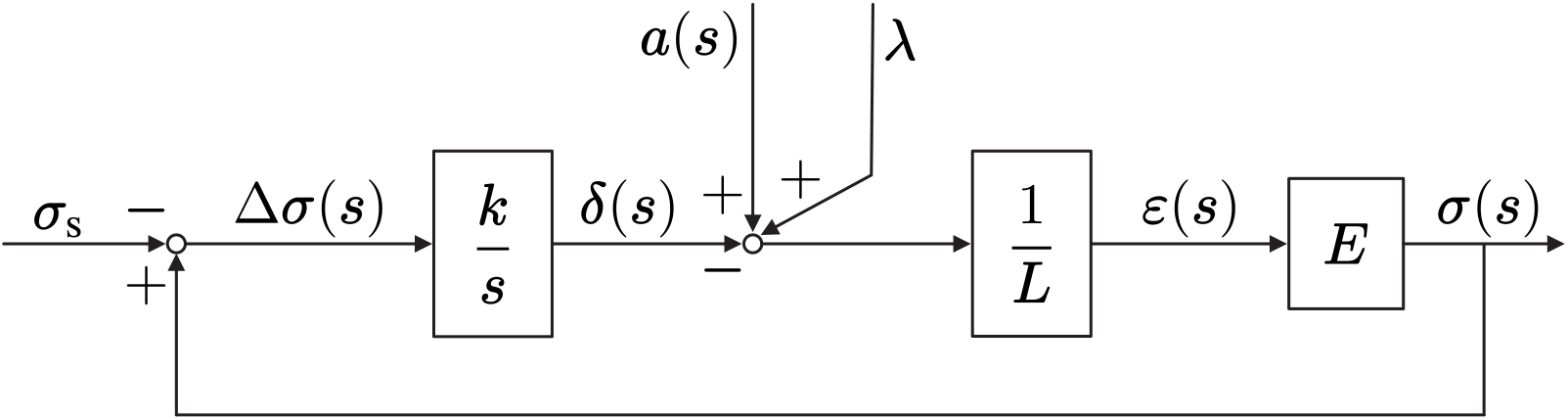
Block diagram of the minimal actin remodeling model. The stress error Δ*σ*(*s*) drives the turnover-mediated rest-length update through an integrator *k/s*. The resulting state *δ*(*s*) enters the elastic relation with a negative sign, forming a negative feedback loop. This structure integrates the stress error and implies complete adaptation under closed-loop stability.

## 4 Closed-loop mathematical structure of mechanics–turnover coupling

### Overview and scope of the formulation

In this section, we reformulate the actin remodeling model in a system-level description to expose a common dynamical structure underlying mechanical adaptation, homeostasis, and remodeling. The purpose of this reformulation is to identify a system structure that enables mechanical adaptation, independent of the specific biomechanical details and functional forms governing mechanical response. By generalizing the actin model at the system level, the formulation allows comparison across scales within a unified description and reveals the integral-feedback structure arising from turnover-mediated remodeling. In the minimal actin model presented in Sec. 2, three features are essential: (i) a mechanically regulated quantity that maintains homeostasis, (ii) a turnover-mediated update of a remodeling state variable, and (iii) feedback coupling whereby updates of the remodeling state modify the mechanical response to environmental disturbances. The general formulation introduced below retains only these system-level architectural elements, while abstracting away the particular molecular mechanisms and the specific forms of the mechanical relation and turnover law.

### General closed-loop formulation via an error representation

#### Definition of variables and governing relations

We introduce three variables: a disturbance input *x*(*t*), a remodeling state variable *y*(*t*), and a homeostatically regulated mechanical quantity *z*(*t*). The regulated quantity is described as a function of the disturbance and the remodeling state variable,

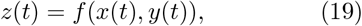

where the specific form of *f* depends on the system under consideration. For example, *f* represents a constitutive stress–strain relation, Laplace’s law relating pressure to wall stress, or the expression for wall shear stress derived from Poiseuille flow. The time evolution of the remodeling state variable *y*(*t*) is governed by a turnover-mediated function *g* of the regulated quantity,

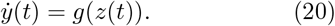

We define the homeostatic setpoint *z*_s_ as an equilibrium of the adaptive dynamics,

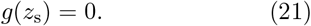

### Error variable and equivalent representation

To express the dynamics explicitly in terms of deviations from homeostasis, we introduce an error variable defined relative to the set-point,

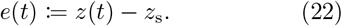

Eq. (20) can then be rewritten as

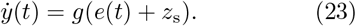

Introducing

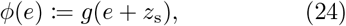

we obtain

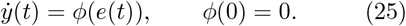

This error representation is a change of variables and does not reduce generality. It makes explicit that turnover-mediated updates are driven by deviations from the set-point (Fig. 5(a)).

**Fig. 5.**
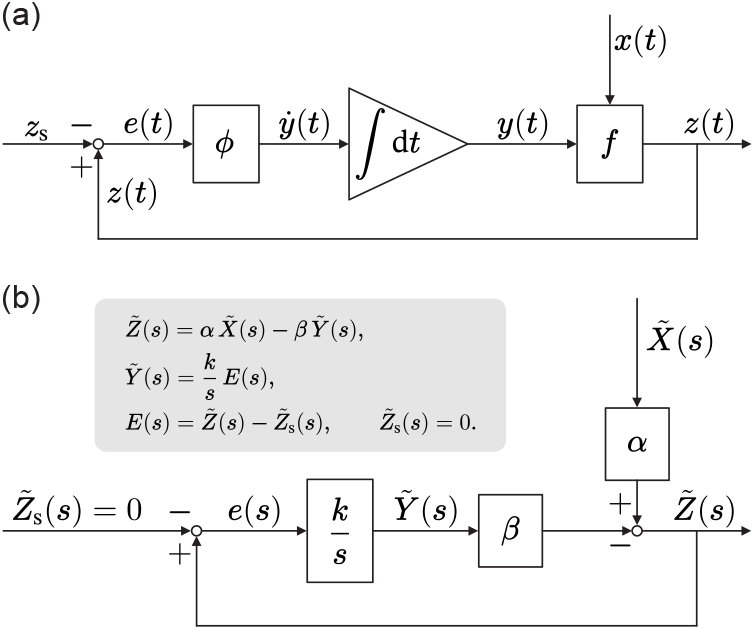
A general formulation and its linearized block-diagram form. (a) A feedback structure with disturbance *x*(*t*), a remodeling state variable *y*(*t*) controlled by turnover, and a homeostatically regulated mechanical quantity *z*(*t*) = *f* (*x*(*t*), *y*(*t*)). The error is defined as *e*(*t*) = *z*(*t*) *− z*_s_ and drives the adaptive dynamics 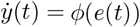, whose time integration yields the remodeling state variable *y*(*t*). (b) Linearization around a homeostatic operating point yields a block-diagram representation with 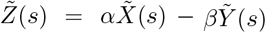 and 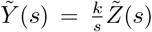. In the linearized setting, the error satisfies 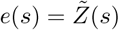.

### Block-diagram interpretation of the adaptive feedback structure

To obtain a block-diagram representation, we linearize the nonlinear formulation around a homeostatic operating point and take its Laplace transform. Let (*x*_0_, *y*_0_) denote a homeostatic operating point. We introduce perturbations

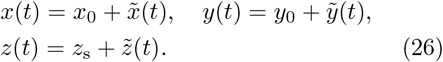

Then

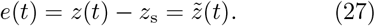

Linearizing the turnover law and the mechanical relation gives

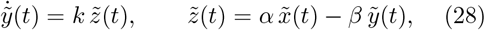

Where

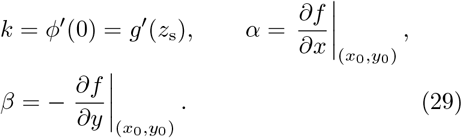

Without loss of generality, the sign of *y* is chosen such that *k >* 0 and *β >* 0. In the Laplace domain, the relations become

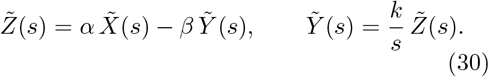

The resulting block diagram is shown in Fig. 5(b), which describes a negative-feedback structure through *β* with integral action through *k/s*. In this linearized form, turnover-mediated state renewal is mathematically equivalent to integral feedback. Under closed-loop stability, this structure yields complete adaptation, i.e., zero steady-state error to step disturbances (Appendix 3).

### Closed-loop mathematical structure of mechanics—turnover coupling in adaptive living systems: *Feedback Adaptive Turnover-mediated Environment-Dependent* system

We refer to this class of turnover-mediated adaptive systems as the *Feedback Adaptive Turnover-mediated Environment-Dependent (FATED)* system class. The name reflects feedback regulation through turnover-mediated structural updates in adaptive living systems (Fig. 6). Here, *Environment-Dependent* indicates that adaptive dynamics are triggered by internal and/or external environmental perturbations acting on the system.

**Fig. 6.**
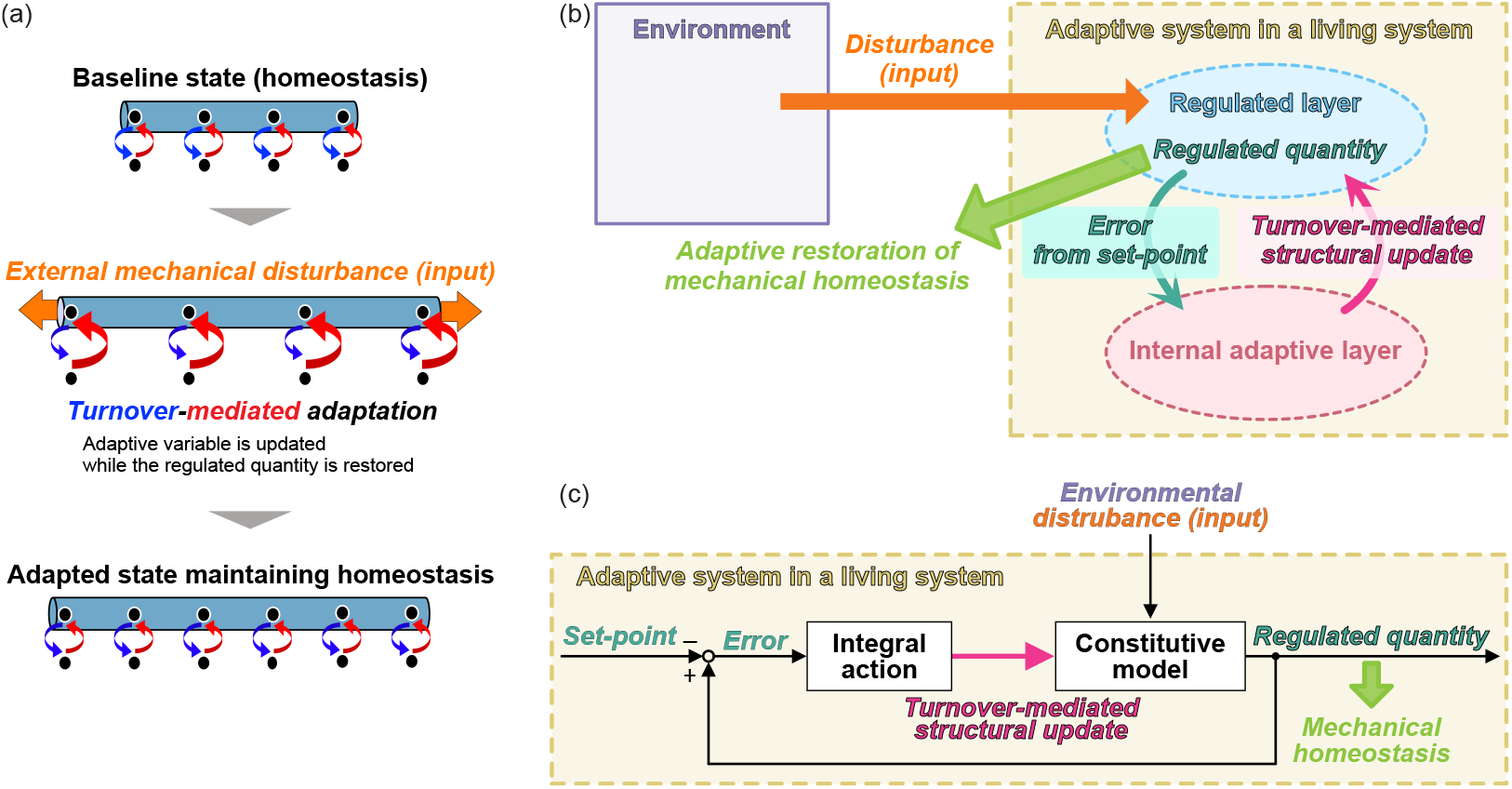
Three equivalent representations of a *FATED* system for mechanical adaptation. (a) Turnover-mediated adaptation at the structural level, illustrating how internal turnover modifies the system while maintaining a regulated mechanical quantity. (b) Conceptual organization of the same mechanism in a living system, highlighting error-driven turnover and adaptive restoration of mechanical homeostasis. (c) Abstract system-level representation of the identical feedback structure, showing how error from a set-point is converted into turnover-mediated structural updates that restore homeostasis.

A defining property of a *FATED* system is that persistent disturbances drive turnover-mediated updates of the remodeling state variable (Eq. (20)), rather than only transient deviations in the regulated quantity. Because the remodeling state variable enters the relation for the regulated quantity (Eq. (19)), turnover can effectively update the internal mechanical state, such as a shift in a reference configuration, a change in effective shape, or material parameters. Thus, adaptation arises from turnover-mediated reconfiguration of the system itself, not from an external controller.

At the system level, the feedback logic can be summarized as follows (Fig. 6). A disturbance perturbs the regulated quantity away from its set-point, and the resulting error modulates turnover. Turnover in turn updates the remodeling state variable *y*(*t*), modifying the disturbance-to-output relation until the error vanishes, at which point adaptation ends. In the linearized canonical representation, the same mechanism appears as a negative-feedback loop with an explicit integrator with gain arising from turnover, which constitutes the minimal structure enabling complete adaptation. Broader examples, such as stress-regulated growth and remodeling of blood vessels or cytoskeletal reorganization under load, can be discussed within the same framework (Rodriguez et al. 1994; Humphrey 2021; Fung and Liu 1989; Humphrey and Schwartz 2021; Hahn and Schwartz 2009). These examples are discussed further in the next section.

## 5 Timescale of turnover-mediated adaptation across scales

In this section, we derive a characteristic timescale relation from the *FATED* structure and use it to compare mechanical adaptation across systems and biological scales, from subcellular to organ levels.

### Characteristic adaptation timescale in the linearized *FATED* system

We start from the linearized common structure of the *FATED* system derived in the previous section (Eq. (28)), with *β >* 0 and *k >* 0. Eliminating 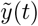 yields a first-order closed-loop dynamics for the regulated quantity,

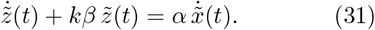

Here, *k* is the local turnover gain, quantifying how strongly deviations of the regulated quantity drive turnover-mediated updates of the remodeling state variable. Similarly, *β* is the local mechanical feedback gain, quantifying how changes in the remodeling state variable affect the regulated quantity. The homogeneous closed-loop dynamics exhibit a single relaxation rate *kβ*, yielding the time constant

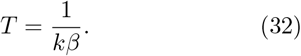

This time constant characterizes the exponential relaxation of the regulated quantity and defines the adaptation time in the linearized regime.

### Turnover-mediated constraint on adaptation time across biological scales

As shown in Eq. (32), the characteristic adaptation time contains the turnover-mediated coefficient *k*. This implies that renewal of the remodeling state variable through turnover is a necessary dynamical mechanism in mechanical adaptation described by the *FATED* system. In principle, direct validation would require independent estimation of *k* and *β* through controlled perturbation experiments. Because such measurements are not systematically available across biological systems, we instead compare reported adaptation times with representative turnover times to assess whether turnover sets the achievable speed of adaptation.

We analyzed representative examples of mechanical adaptation spanning subcellular, cellular, tissue, and organ scales (Table 2). For each system, we identified the three *FATED* variables: an environmental mechanical disturbance *x*, a remodeling state variable *y* updated through turnover, remodeling, or growth, and (iii) a regulated mechanical quantity *z* maintained near a homeostatic set-point. From reported experiments or phenomenological models, we extracted an adaptation time *T*_ad_ characterizing the relaxation of *z*, and a corresponding turnover time *T*_turn_ defined as a representative turnover timescale for the structural process. The operational definitions of *T*_turn_ and *T*_ad_ for each system are summarized in Appendix 4.

**Table 2.** Mechanical adaptation systems analyzed in Fig. 7. For each system, the disturbance *x*, remodeling state variable *y*, and regulated mechanical quantity *z* are identified within the *FATED* system, together with the corresponding turnover time *T*_turn_ and adaptation time *T*_ad_ (reported ranges).

| No. | System | $x$ | $y$ | $z$ | $T_{\text{turn}}$ | $T_{\text{ad}}$ | Refs. |
| --- | --- | --- | --- | --- | --- | --- | --- |
| 1 | Focal adhesion stress homeostasis | Applied force or stiffness change | Adhesion size and composition | Adhesion traction stress | 4–17 min | 1–10 min | (Balaban et al. 2001; Riveline et al. 2001; Kubow et al. 2017; Schwingel and Bastmeyer 2013; Stehbens and Wittmann 2014) |
| 2 | Microtubule lattice self-repair | Lattice damage | Tubulin incorporation | Microtubule stiffness | 1–2 min | 1–2 min | (Schaedel et al. 2015; Aumeier et al. 2016; Portran et al. 2017) |
| 3 | Yeast plasma membrane tension regulation | Osmolarity decrease | Membrane area reservoir | Membrane tension | 2–6 min | 2–6 min | (Lemière et al. 2021; Kabeche et al. 2015) |
| 4 | Division-mediated epithelial stress relaxation | Uniaxial stretch | Division orientation | Monolayer stress | 50–60 min | 50 min–3 h | (Wyatt et al. 2015) |
| 5 | Flow-mediated arterial caliber remodeling | Flow rate change | Vessel inner diameter | Wall shear stress | 1–7 d | 2–4 wk | (Langille and O’Donnell 1986; Cho et al. 1997; Sho et al. 2001; Tronc et al. 2000; Nuki et al. 2009; Bakker et al. 2008) |
| 6 | Pressure-mediated arterial wall remodeling | Intraluminal pressure increase | Wall thickness | Circumferential wall stress | 3–14 d | 2–12 wk | (Owens and Reidy 1985; Hu et al. 2008; Kuang et al. 2013; Matsumoto and Hayashi 1994; Wolinsky 1970) |
| 7 | Poplar stem gravitropic righting | Gravity direction change | Asymmetric radial growth | Stem tilt | 7–14 d | 2–6 wk | (Coutand et al. 2007; Liu et al. 2021) |
| 8 | Pressure overload cardiac wall remodeling | Afterload increase | Wall thickness | Circumferential wall stress | 3–14 d | 2–6 wk | (Martin et al. 1977; Everett et al. 1977; Liao et al. 2002; Wang et al. 2023; Zi et al. 2019) |
| 9 | Bone mechanostat adaptation | Cyclic mechanical loading | Surface bone formation | Strain level | 5–14 d | 7–21 d | (Sun et al. 2017; McBride and Silva 2012; Yang et al. 2020) |

Across systems, *T*_ad_ is generally comparable to or longer than *T*_turn_ (Fig. 7). This observation is consistent with the linear *FATED* system Eq. (32), in which the adaptation time depends on the product *kβ*, where *k* reflects the local gain of turnover-mediated remodeling and *β* the mechanical coupling from remodeling to the regulated quantity. Because *k* and *β* are generally system-dependent and are not directly measurable in most cases, we do not expect a one-to-one quantitative relation between *T*_ad_ and *T*_turn_. Instead, the cross-scale trend suggests that turnover-mediated renewal introduces a fundamental timescale into the adaptive loop and can act as a practical constraint on how fast mechanical adaptation can proceed.

**Fig. 7.**
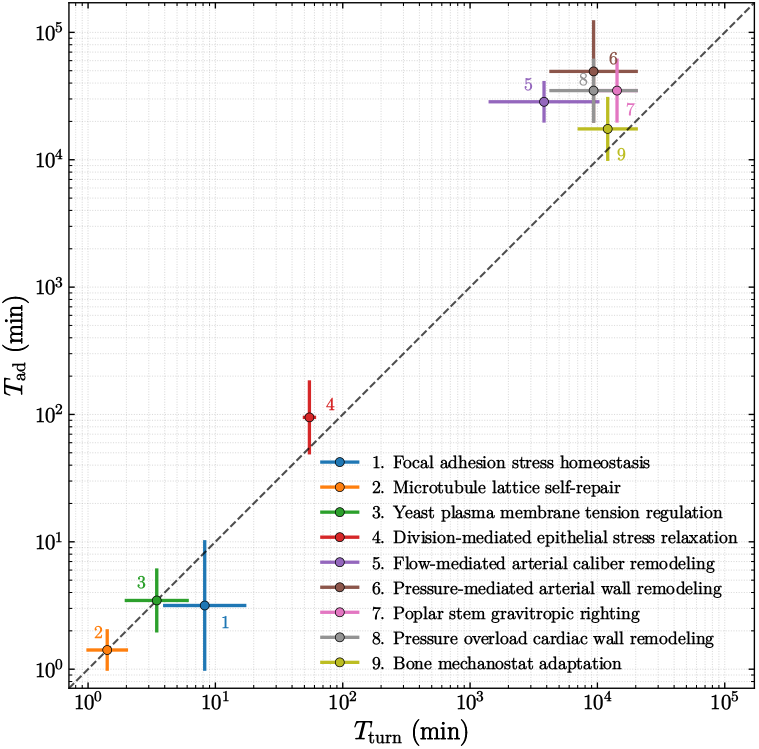
Comparison of adaptation and turnover timescales across biological systems. Log–log plot of the adaptation time *T*_ad_ versus the turnover time *T*_turn_ for the representative systems listed in Table 2. The dashed line indicates *T*_ad_ = *T*_turn_. Across systems, adaptation times are typically on the order of or longer than turnover times. This trend supports the interpretation that turnover-mediated renewal introduces a fundamental constraint on the achievable rate of mechanical adaptation, consistent with the time constant predicted by the *FATED* system.

## 6 Discussion

In this study, we describe the minimal mathematical structure of closed-loop mechanics–turnover coupling that ensures turnover-mediated homeostasis and intrinsically includes integral action, providing a unified understanding of mechanically adaptive remodeling across scales. Based on in vitro experiments, we formulate a remodeling model of mechanically regulated actin turnover, showing how turnover-mediated updating of the rest length restores mechanical stress under sustained deformation (Sec. 2). This stress–turnover feedback reveals an integral-action mechanism in the closed-loop dynamics (Sec. 3). Generalizing beyond actin, we isolate the closed-loop mathematical structure in terms of a disturbance input, a homeostatically regulated mechanical quantity, and a remodeling state variable renewed by turnover. Using this framework, we show how structural renewal feeds back onto mechanics (Sec. 4). This structure yields a characteristic adaptation time constant set by the effective strength of mechanics–turnover coupling (Sec. 5). We term this structural class *Feedback Adaptive Turnover-mediated Environment-Dependent (FATED)* systems. Finally, we assess cross-system consistency by comparing reported adaptation and turnover timescales across representative remodeling systems (Fig. 7, Table 2), enabling direct comparison of mechanically adaptive remodeling across scales within the *FATED* framework (Sec. 5).

We first consider *in vitro* actin reconstitution experiments in which mechanical loading modulates filament turnover to promote elongation (Fig. 1). Mechanically, loading increases the stress supported by the filament, while turnover-mediated elongation can be interpreted as an increase in the rest length. This coupling drives the stress back toward its homeostatic value. This interpretation motivates the remodeling model, in which stress-regulated turnover is coarse-grained as rest-length updating coupled to the elastic stress–strain relation under an imposed deformation (Sec. 2). This model shows how stress-regulated turnover can generate mechanical adaptation with a characteristic timescale (Fig. 3).

The central feature of the *FATED* system is that the deviation of a homeostatically regulated mechanical quantity from its reference level regulates the turnover rate, which updates the remodeling state variable. In the linearized *FATED* system, turnover-mediated remodeling is mathematically equivalent to integral action, providing a basis for adaptation. This conclusion is not limited to linear elastic materials, but also applies to viscoelastic or nonlinear mechanical elements (Appendices 5 and 6).

To consider why turnover is indispensable for mechanical adaptation, we note that biological structures are continuously renewable. This renewability stems from modular assemblies stabilized by non-covalent interactions, which allow constituent exchange while preserving function (Alberts et al. 2022; Misteli 2001). This exchange-ability enables continuous replacement and reorganization of the components; thus, turnover provides a generic route for updating the internal structure. Adaptive living systems share this turnover-mediated renewability as captured by *FATED* systems.

To connect the structural prediction to observations across scales, we assembled representative examples of mechanical adaptation and defined two operational timescales for each system: the adaptation time and the turnover time (Table 2 and Appendix 3). Across representative systems, the adaptation time and turnover time are of the same order, with the turnover time typically no longer than the adaptation time (Fig. 6). We do not claim causality. This is a consistency check. If turnover drives adaptation, then renewal should be at least as fast as the observed adaptation.

The interpretation proposed here for *FATED* systems relates to, but is distinct from, several existing theoretical approaches. In systems biology, integral feedback has been widely studied as a core mechanism enabling perfect adaptation in biochemical reaction networks. Studies on bacterial chemotaxis, calcium homeostasis, and antithetic integral feedback controllers have shown that robust elimination of steady-state error typically requires an internal integral action within the regulatory network (Yi et al. 2000; Briat et al. 2016; Aoki et al. 2019). However, these studies primarily concern biochemical concentration dynamics and reaction-network motifs. In contrast, our approach focuses on mechanically adaptive remodeling at the system level. In biomechanics, stress-dependent growth and remodeling models have long incorporated constitutive laws in which structural change is driven by deviations from a reference stress (Rodriguez et al. 1994; Humphrey 2021; Ambrosi and Guana 2007). While these formulations capture phenomenological stress-regulated structural updating, they are typically formulated as constitutive relations within continuum mechanics. Our framework formalizes a closed-loop mathematical structure based on concepts from control theory, showing that turnover provides an intrinsic integral action across biological hierarchies. In classical artificial control systems, integral action is implemented through a dedicated controller state that is externally designed and coupled to a plant via a control input (Åström and Murray 2021). In *FATED* systems, by contrast, no external controller is imposed, while integral-equivalent action emerges intrinsically.

Controlled perturbation experiments can directly identify the two local gains that set the adaptation rate: the turnover gain *k* and the mechanical feedback gain *β*. With *k* and *β* measured, the predicted timescale in Eq. (32) becomes a quantitative, testable link between theory and experiment, enabling direct comparison across systems. Finally, many other mechanically adaptive systems involve multiple coupled variables and hierarchical organization across scales, motivating extensions of the framework to multi-variable and multi-layer *FATED* systems.

## Appendix 1: Ramp response in the actin remodeling model

### Model and role of linearization

We consider the fixed-end actin filament model used in the main text, where the filament experiences an imposed deformation *a*(*t*) and a turnover-driven change of the rest length *δ*(*t*). In this Supplement, numerical plots are obtained by directly solving the full nonlinear model. In parallel, we introduce a small-deformation linearization solely to derive analytic expressions and to identify the characteristic timescale governing the transient dynamics.

### Linearized dynamics (Hookean elasticity)

Under the standard small-deformation assumption *L* ≫ *λ, a*(*t*), *δ*(*t*), the strain increment is approximated as

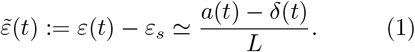

For a Hookean solid with modulus *E*, the stress increment becomes

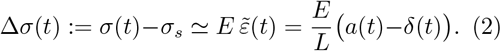

Turnover is modeled by the mechano-chemical feedback

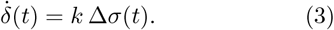

Combining Eqs. (2) and (3) yields the closed-loop linear dynamics

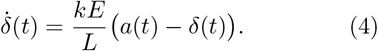

This linearization exposes a single characteristic adaptation timescale

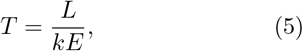

which governs the exponential relaxation in the Hookean limit.

### Analytic ramp response (linearized)

We apply a ramp deformation

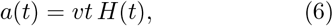

where *v* is the ramp rate and *H*(*t*) is the Heaviside step function.

Solving Eq. (4) with *δ*(0) = 0 gives

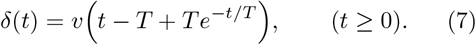

The stress increment follows from Eq. (2):

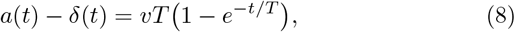

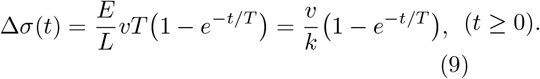

### Parameters

Unless otherwise stated, we use the same parameter values as in the main-text step-response figure (Table 1). For the ramp input, we set the ramp rate to *v* = 0.6 nm*/*s.

### Interpretation

The linearized ramp response exhibits a nonzero steady-state stress offset:

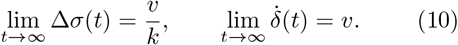

Thus, under sustained ramp deformation, the turnover-driven rest-length change asymptotically matches the ramp rate 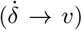, while the stress error converges to the constant Δ*σ*_∞_ = *v/k*. Importantly, this steady offset is independent of the elastic modulus *E* and the filament length *L*; it depends only on the ramp rate *v* and the mechanochemical sensitivity *k*. The transient approach to this regime is controlled by the characteristic timescale *T* = *L/*(*kE*) identified above. These behaviors are confirmed by numerical solutions of the full nonlinear model (Fig. A1), and the steady offset is further verified by the collapse of the normalized stress error *k*(*σ−σ*_*s*_)*/v* → 1 (Fig. A2).

**Fig. A1.**
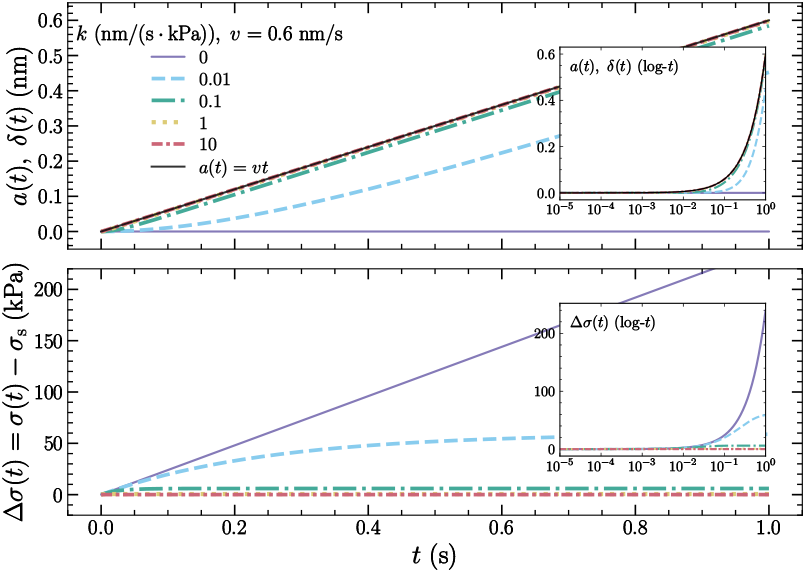
Ramp response in the actin remodeling model (numerical solution of the full nonlinear model). The imposed deformation is a ramp *a*(*t*) = *vt* with *v* = 0.6 nm*/*s. **(Top)** *a*(*t*) and the turnover-driven change of rest length *δ*(*t*) for different mechano-chemical sensitivities *k*. **(Bottom)** stress error Δ*σ*(*t*) = *σ*(*t*) *− σ*_*s*_. For *k >* 0, Δ*σ*(*t*) approaches a constant plateau, consistent with the steady offset Δ*σ*_∞_ = *v/k* predicted by the linearized analysis. Insets show the same quantities on a logarithmic time axis to highlight early-time transients.

**Fig. A2.**
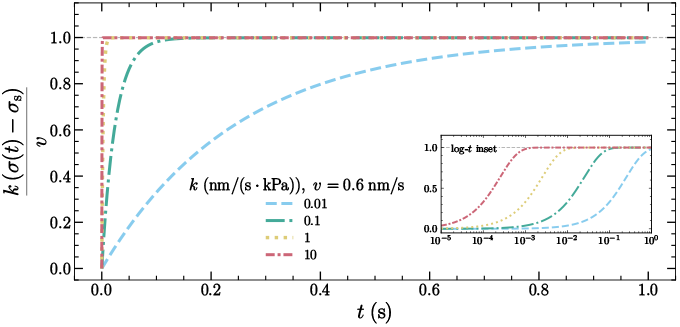
Dimensionless collapse of the ramp response. Using the same numerical solutions as Fig. A1, we plot the normalized stress error *k*(*σ*(*t*) − *σ*_*s*_)*/v* for *k >* 0 (the case *k* = 0 is omitted because the normalization is not informative). All curves converge to unity, demonstrating Δ*σ*_∞_ = *v/k* without drawing multiple horizontal guides. The inset shows early-time dynamics on a logarithmic time axis.

## Appendix 2: Proof of complete adaptation in the actin remodeling model

Here, we provide a formal proof of complete adaptation in the linearized actin remodeling model using the final value theorem. The proof clarifies how turnover-mediated rest-length adaptation guarantees zero steady-state stress error in response to step-like mechanical perturbations.

### Definition of complete adaptation

We define *complete adaptation* as the vanishing of the steady-state error of the regulated mechanical variable. In the present context, this corresponds to zero steady-state stress error,

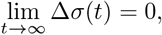

in response to a step deformation input.

### Linearized dynamics

We consider the linearized actin remodeling model. Defining the stress error as

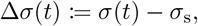

the turnover law linearizes to

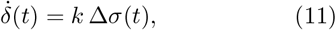

where *δ*(*t*) denotes the adaptive change in the rest length and *k >* 0 is the turnover gain.

Let *a*(*t*) denote the externally imposed deformation, and assume zero initial conditions. Taking the Laplace transform of Eq. (11) yields

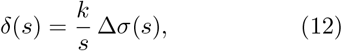

which corresponds to an integrator with gain *k*.

### Closed-loop stress response

In the linearized kinematics, the elastic strain is given by

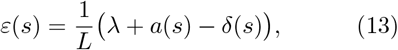

where *L* is the reference rest length and *λ* is a constant baseline deformation. The constitutive relation relates stress and strain as

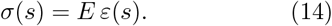

The baseline deformation *λ* sets the homeostatic stress *σ*_s_ = *Eλ/L* and cancels out when forming the stress error Δ*σ*(*s*) = *σ*(*s*) − *σ*_s_.

Substituting Eq. (12)–Eq. (14) gives the closed-loop relation

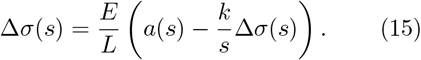

Solving for the stress response yields the closedloop transfer function

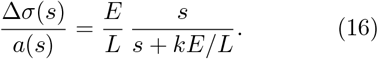

### Final value theorem and complete adaptation

The transfer function Eq. (16) has a single pole at *s* = *−kE/L*, indicating closed-loop stability for *k >* 0. For a step deformation input *a*(*t*) = *a*_0_*H*(*t*) with *a*(*s*) = *a*_0_*/s*, the final value theorem gives

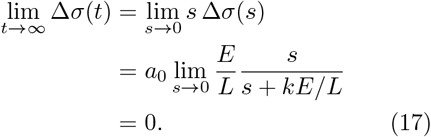

Thus, the linearized actin remodeling model exhibits complete adaptation: the steady-state stress error vanishes in response to step-like deformations. This result follows directly from the integral action introduced by turnover-mediated rest-length adaptation and is independent of molecular details.

## Appendix 3: Proof of complete adaptation in the general linear FATED model

Here, we provide a formal proof of complete adaptation in the general linear FATED model using the final value theorem. This proof applies to the linearized common structure and is independent of system-specific molecular details.

We define complete adaptation as zero steady-state error of the regulated quantity in response to a step disturbance. Specifically, we consider a step disturbance

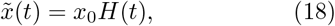

whose Laplace transform is 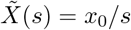.

The linearized common model is given by

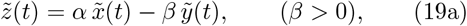

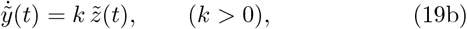

where 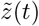 denotes the deviation of the regulated quantity from its set-point. Taking the Laplace transform with zero initial conditions yields

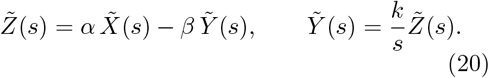

Eliminating 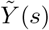 gives the closed-loop transfer function from disturbance to regulated quantity,

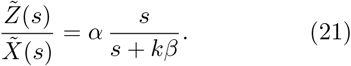

This transfer function has a single pole at *s* = *−kβ*; thus, the closed loop is asymptotically stable for *kβ >* 0, and the final value theorem applies.

Applying the final value theorem, we obtain

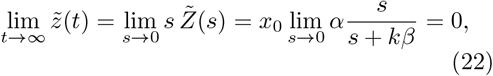

provided that the closed-loop system is stable.

This result demonstrates that the embedded integrator in the adaptive dynamics is sufficient to guarantee complete adaptation, i.e., zero steady-state error to step disturbances, for the entire class of systems described by the linear FATED structure.

## Appendix 4: Operational definitions of turnover and adaptation timescales for Fig. 6

**Table A1.** Operational definitions for timescales. For each system, we specify how the turnover time *T*_turn_ and the adaptation time *T*_ad_ were operationally defined from the cited sources. Ranges reflect reported variability across experimental conditions or measurement protocols, and definitions emphasize the renewal process most directly responsible for updating the adaptive variable in each system.

| No. | System | $T_{\text{turn}}$ | $T_{\text{ad}}$ | Operational definition |
| --- | --- | --- | --- | --- |
| 1 | Focal adhesion stress homeostasis | 4–17 min | 1–10 min | **Turnover:** Turnover time is taken as the lifetime of individual focal adhesion assemblies measured by live imaging. (Kubow et al. 2017; Stehbens and Wittmann 2014)<br>**Adaptation:** Adaptation time is defined as the minutes-scale stabilization of adhesion traction stress after mechanical perturbation, quantified over a $\sim 1$–10 min window. (Riveline et al. 2001; Schwingel and Bastmeyer 2013) |
| 2 | Microtubule lattice self-repair | 1–2 min | 1–2 min | **Turnover:** Turnover time is defined as the time required for tubulin incorporation into mechanically damaged microtubule lattices, visualized by incorporation of fluorescently labeled tubulin at damage sites after laser-induced lattice damage. (Schaedel et al. 2015)<br>**Adaptation:** Adaptation time corresponds to recovery of stiffness, quantified by the return of persistent length during rest after cyclic bending. (Schaedel et al. 2015) |
| 3 | Yeast plasma membrane tension regulation | 2–6 min | 2–6 min | **Turnover:** Turnover reflects membrane-area renewal after hypotonic shock, occurring over the 2–6 min window during which endocytosis transiently decreases and subsequently recovers while exocytosis is enhanced. (Lemière et al. 2021; Kabeche et al. 2015)<br>**Adaptation:** Adaptation corresponds to restoration of membrane tension, quantified by recovery of endocytic density within 2–6 min after osmotic perturbation. (Lemière et al. 2021) |
| 4 | Division-mediated epithelial stress relaxation | 50–60 min | 50 min–3 h | **Turnover:** Turnover time is defined as the duration from mitotic entry following uniaxial stretch to daughter-cell reintegration into the monolayer. (Wyatt et al. 2015)<br>**Adaptation:** Adaptation time spans from the onset of division-associated remodeling at approximately 50 min to the subsequent hours-scale evolution of monolayer stress after stretch. (Wyatt et al. 2015) |
| 5 | Flow-mediated arterial caliber remodeling | 1–7 d | 2–4 wk | **Turnover:** Turnover time is read from early remodeling-marker time courses after flow alteration, including apoptosis and matrix metalloproteinase activation peaking within several days. (Cho et al. 1997; Sho et al. 2001; Tronc et al. 2000)<br>**Adaptation:** Adaptation time is determined from longitudinal measurements of lumen diameter, which progressively decreases or increases over 2–4 wk following sustained flow rate change. (Langille and O'Donnell 1986; Nuki et al. 2009) |
| 6 | Pressure-mediated arterial wall remodeling | 3–14 d | 2–12 wk | **Turnover:** Turnover time is read from early time courses of medial smooth muscle proliferation and matrix synthesis following pressure elevation, detected within 3–14 d. (Owens and Reidy 1985; Hu et al. 2008)<br>**Adaptation:** Adaptation time is determined from serial measurements of arterial wall thickness or medial area, which increase progressively over 2–12 wk under sustained pressure overload. (Kuang et al. 2013; Matsumoto and Hayashi 1994; Wolinsky 1970) |
| 7 | Poplar stem gravitropic righting | 7–14 d | 2–6 wk | **Turnover:** Turnover time is inferred from the lag between stem reorientation and measurable asymmetric radial growth leading to tension wood formation, observed over 7–14 d. (Coutand et al. 2007)<br>**Adaptation:** Adaptation time is determined from longitudinal measurements of stem curvature and tilt, which progressively reorient toward the vertical over 2–6 wk following a change in gravity direction. (Coutand et al. 2007) |
| 8 | Pressure overload cardiac wall remodeling | 3–14 d | 2–6 wk | **Turnover:** Turnover time is inferred from early time-course measurements of ventricular hypertrophy, including increases in heart weight within 3–14 d after pressure overload. (Everett et al. 1977)<br>**Adaptation:** Adaptation time is determined from longitudinal measurements of ventricular wall thickness in transverse aortic constriction models, which show progressive thickening over 2–6 wk under sustained afterload elevation. (Liao et al. 2002; Wang et al. 2023; Zi et al. 2019) |
| 9 | Bone mechanostat adaptation | 5–14 d | 7–21 d | **Turnover:** Turnover time is taken as the delay from cyclic loading to the onset of lamellar surface bone formation, read from dynamic histomorphometry in axial tibial loading experiments. (Sun et al. 2017; McBride and Silva 2012)<br>**Adaptation:** Adaptation time is inferred from the time window over which cortical structure significantly increases after cyclic loading, based on longitudinal microCT and time-resolved morphometry over 7–21 d. (Yang et al. 2020; Sun et al. 2017) |

## Appendix 5: Linear viscoelastic remodeling model

### Motivation and role of linearization

In the main text, we introduced a minimal remodeling model in which mechanically regulated turnover updates the rest length *δ*(*t*) under a fixed-end deformation protocol. While the minimal version assumes purely elastic filament mechanics, actin filaments have also been reported to exhibit viscous mechanical responses (Shamloo and Mehrafrooz 2018b; Sato et al. 1985; Zaner and Stossel 1982; Kojima et al. 1994), and many subcellular, cellular, and tissue structures in living systems are more naturally described as viscoelastic materials (Fletcher and Mullins 2010; Bao and Suresh 2003; Chaudhuri et al. 2020). Motivated by this broader biomechanical context, here we generalize the remodeling framework to a standard linear solid (SLS), a canonical three-parameter linear viscoelastic model capturing a finite equilibrium modulus and a characteristic relaxation time.

In this Supplement, numerical plots are obtained by directly solving the full nonlinear kinematics, as in the main text. In parallel, we introduce a small-deformation linearization solely to obtain analytic expressions and to identify the characteristic timescales that control transient dynamics. This separation is intentional: linearization is used as a diagnostic tool for timescales and asymptotic limits, whereas numerical simulations retain the nonlinear kinematics.

### SLS constitutive law and linearized kinematics

We model the filament rheology as a standard linear solid (SLS) with elastic moduli *E*_1_, *E*_2_ and viscosity *η*. The constitutive equation is

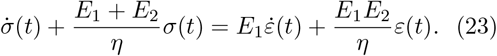

Under the small-deformation assumption *L*≫ *λ, a*(*t*), *δ*(*t*), the strain increment is approximated by

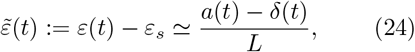

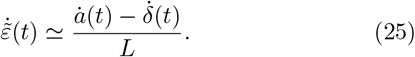

We define the stress increment Δ*σ*(*t*) := *σ*(*t*) *− σ*_*s*_.

Turnover is modeled by the mechano-chemical feedback

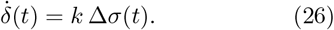

### Closed-loop linear dynamics and characteristic timescales

Subtracting the steady state from Eq. (23) and using Eqs. (25)–(26) yields a second-order linear ODE for *δ*(*t*):

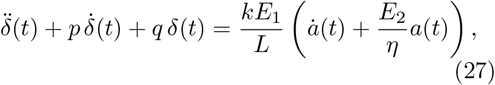

With

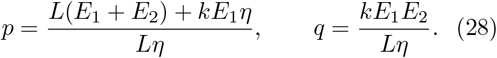

The homogeneous dynamics are governed by the two roots *α*_1,2_ of *s*^2^ + *ps* + *q* = 0:

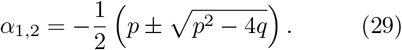

When *p*^2^ *>* 4*q*, the transient response contains two exponential modes 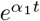 and 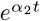, giving two characteristic timescales

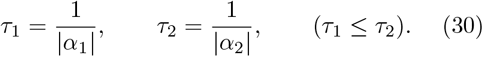

In practice, the late-time approach is dominated by the slower mode *τ*_2_.

### Analytic step response (linearized)

We apply a step deformation *a*(*t*) = *a*_*s*_*H*(*t*). Because Eq. (27) contains 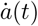, an instantaneous step implies an impulse 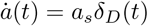 and therefore a jump in 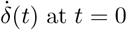 at *t* = 0. Assuming the system is at rest for *t <* 0, *δ*(0^*−*^) = 0 and 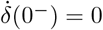, integrating Eq. (27) across *t* = 0 yields 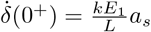.

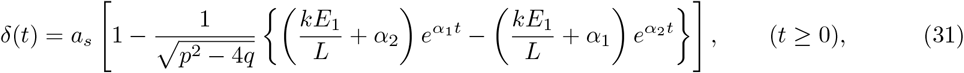

and the stress increment follows from 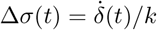:

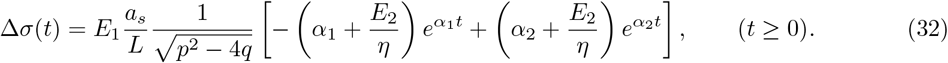

### Analytic ramp response (linearized)

We apply a ramp deformation *a*(*t*) = *vt H*(*t*). Solving Eq. (27) gives

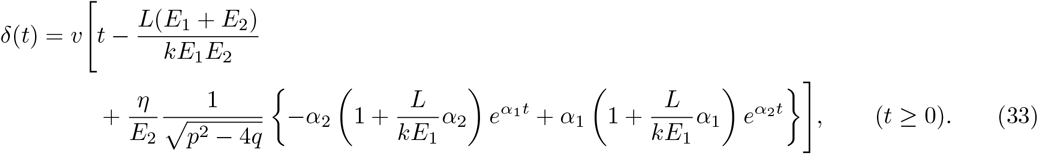

Using 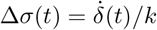, we obtain

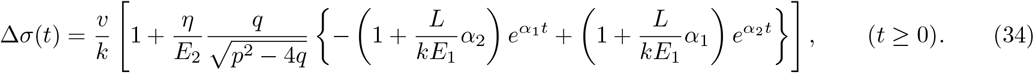

### Interpretation

For the step input, remodeling drives Δ*σ*(*t*) → 0 as *t* → ∞, consistent with perfect adaptation to a constant deformation; viscoelasticity affects only the transient relaxation through the timescales *τ*_1,2_. For the ramp input, the system approaches a steady regime where the rest length tracks the ramp rate and the stress error converges to a nonzero constant:

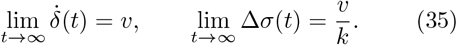

Notably, the steady offset Δ*σ*_∞_ = *v/k* is independent of *E*_1_, *E*_2_, and *η*; viscoelasticity affects only the transient dynamics through the timescales *τ*_1,2_. These behaviors are confirmed by numerical solutions of the full nonlinear model for the step input (Fig. A3) and the ramp input (Fig. A4). The viscosity-independence of the steady ramp offset is further verified by the dimensionless collapse (Fig. A5) and by sweeping *η* over multiple decades (Fig. A6).

## Appendix 6: Nonlinear elastic remodeling model (strain-stiffening)

### Motivation

The minimal remodeling model in the main text assumes a linear elastic constitutive law. Here, we introduce a minimal nonlinear elastic generalization with strain-stiffening. The purpose of this Supplement is threefold: (i) to show that the step-induced stress jump depends nonlinearly on the deformation amplitude, (ii) to clarify that the steady adapted state under a step input is still determined purely by fixed-end kinematics, and (iii) to demonstrate that the late-time relaxation can be characterized by a local linearization governed by the tangent modulus.

**Fig. A3.**
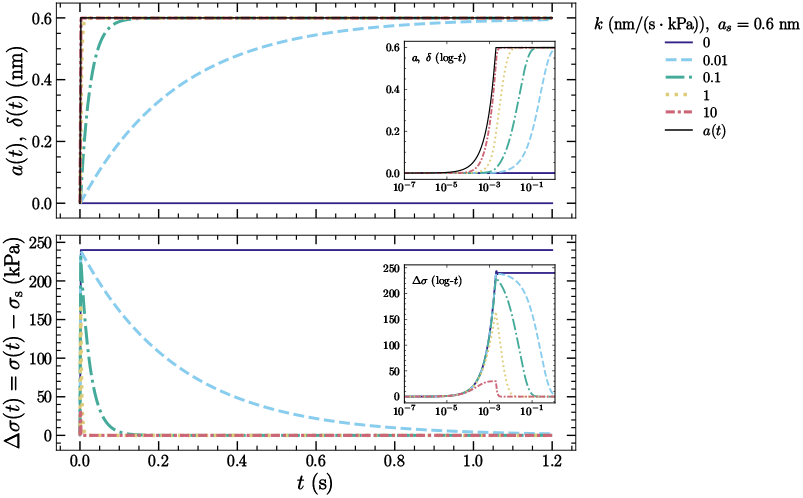
Step response in the SLS-based remodeling model (numerical solution with full nonlinear kinematics). The imposed deformation is a step of amplitude *a*_*s*_ = 0.6 nm, implemented as a smooth step with rise time *t*_rise_ = 2.0 × 10^*−*3^ s for numerical stability. Model parameters are *L* = 5.0 *µ*m, *λ* = 5.0 nm, *E*_1_ = *E*_2_ = 4.0 GPa, *η* = 5.0 Pa s, and *k* = 0, 0.01, 0.1, 1, 10 nm*/*(s · kPa). **(Top)** Input *a*(*t*) and rest-length change *δ*(*t*). **(Bottom)** Stress error Δ*σ*(*t*) = *σ*(*t*) *− σ*_*s*_. For *k >* 0, remodeling drives *δ*(*t*) → *a*_*s*_ and relaxes the stress error toward Δ*σ*(*t*) → 0 (perfect adaptation to a constant deformation), with the transient shaped by the SLS relaxation and the remodeling feedback. For *k* = 0, no remodeling occurs and the stress error remains elevated. Insets show early-time dynamics on a logarithmic time axis.

### Nonlinear constitutive law and nonlinear kinematics

We keep the fixed-end kinematics and turnover law as in the main text. The strain is computed from the full nonlinear kinematics

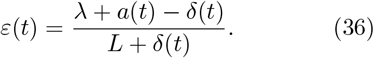

We adopt a minimal nonlinear elastic constitutive law (cubic strain-stiffening)

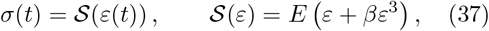

where *E* is the small-strain modulus and *β >* 0 controls strain-stiffening.

The homeostatic reference state is defined by

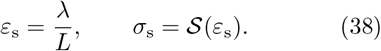

**Fig. A4.**
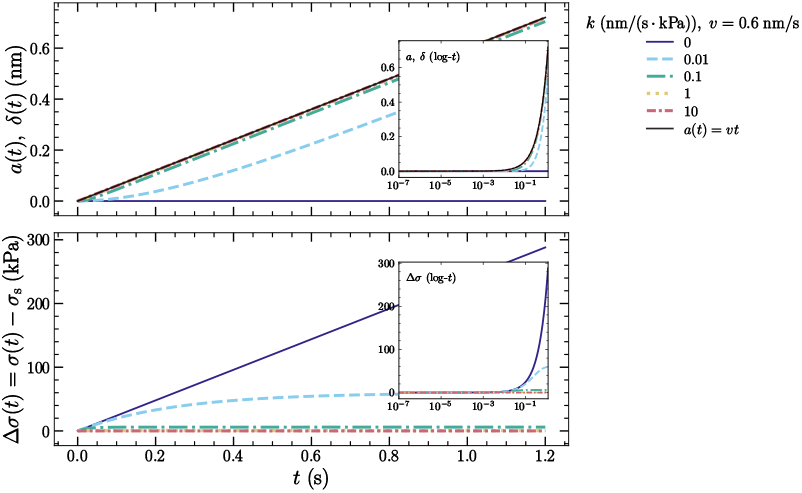
Ramp response in the SLS-based remodeling model (numerical solution with full nonlinear kinematics). The imposed deformation is a ramp *a*(*t*) = *vt* with *v* = 0.6 nm*/*s. Filament rheology is modeled as a standard linear solid with *E*_1_ = *E*_2_ = 4.0 GPa and *η* = 5.0 Pa s, and remodeling is driven by 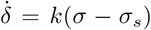 with *k* = 0, 0.01, 0.1, 1, 10 nm*/*(s · kPa). **(Top)** Input *a*(*t*) and turnover-driven rest-length change *δ*(*t*). **(Bottom)** Stress error Δ*σ*(*t*) = *σ*(*t*) − *σ*_*s*_. For *k >* 0, Δ*σ*(*t*) approaches a constant plateau, consistent with the steady offset Δ*σ*_∞_ = *v/k* predicted by the linearized analysis, while 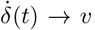. Insets show the same quantities on a logarithmic time axis to highlight early-time transients.

**Fig. A5.**
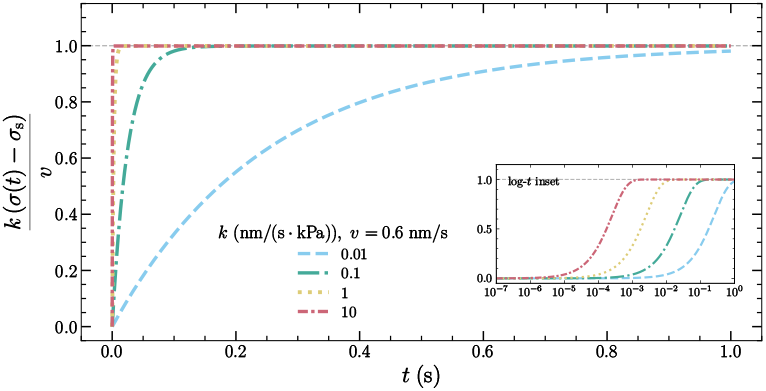
Dimensionless verification of the ramp steady offset in the SLS-based remodeling model. Using the same ramp simulations as Fig. A4, we plot the normalized stress error *k*(*σ*(*t*) − *σ*_*s*_)*/v* for *k >* 0 (the case *k* = 0 is omitted because the normalization is not informative). All curves converge to unity, providing a parameter-independent confirmation of Δ*σ*_∞_ = *v/k* without adding multiple horizontal guides in Fig. A4. The inset shows early-time dynamics on a logarithmic time axis.

**Fig. A6.**
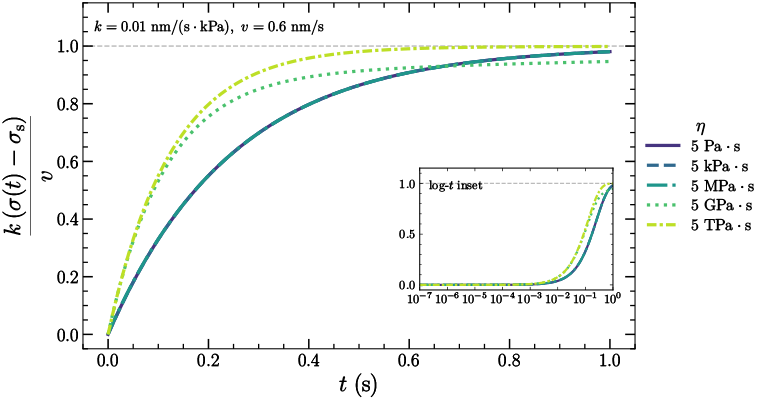
Viscosity sweep of the ramp response in the SLS-based remodeling model (dimensionless verification). The imposed deformation is a ramp *a*(*t*) = *vt* with *v* = 0.6 nm*/*s. Filament rheology is modeled as a standard linear solid with *E*_1_ = *E*_2_ = 4.0 GPa and viscosity *η*, and remodeling is driven by 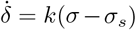 with *k* = 0.01 nm*/*(s · kPa). Other parameters are *L* = 5.0 *µ*m and *λ* = 5.0 nm. We sweep viscosity as *η* = *η*_0_ × 10^*m*^ with *η*_0_ = 5.0 Pa s and *m* ∈ {0, 3, 6, 9, 12}, and report *η* using engineering prefixes for readability (Pa s, kPa s, MPa s, GPa s as appropriate). Curves show the normalized stress error *k*(*σ*(*t*) − *σ*_*s*_)*/v*, which converges to unity for all *η*, confirming the viscosity-independence of the steady ramp offset Δ*σ*_∞_ = *v/k*. Viscosity affects only the transient approach to this limit. The inset shows the same quantity on a logarithmic time axis to highlight early-time transients.

Turnover is modeled by the same mechano-chemical feedback as in the main text,

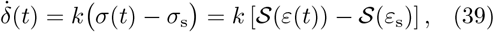

with *δ*(0) = 0 and *a*(0) = 0.

### Step input and amplitude-dependent stress jump

We apply a step deformation *a*(*t*) = *a*_*s*_*H*(*t*). Since *δ*(0) = 0, we have *ε*(0^+^) = (*λ* + *a*_*s*_)*/L* = *ε*_s_ + *a*_*s*_*/L*.

Because *δ*(*t*) is continuous, the strain and stress exhibit an instantaneous jump at *t* = 0:

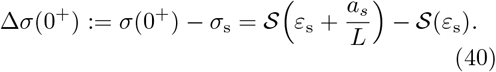

For the cubic constitutive law Eq. (37), this becomes

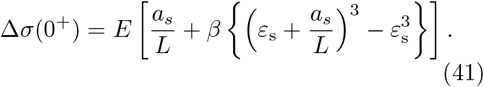

(41)

For sufficiently small *a*_*s*_*/L*, the jump reduces to a local linear form,

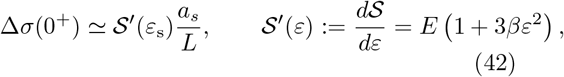

showing that the apparent stiffness for small perturbations is given by the tangent modulus at the homeostatic strain.

### Steady state under step input

At steady state, 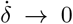 implies *σ* → *σ*_s_. For a monotone constitutive law *S* (as in Eq. (37) with *β >* 0), this yields *ε* → *ε*_s_ uniquely. Therefore, the steady rest-length change is determined purely by kinematics:

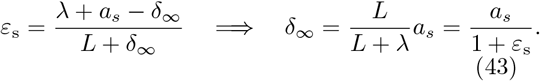

Thus, under a step deformation, the steady outcome is independent of the constitutive parameters (*E, β*).

### Local linearization and effective relaxation time

To characterize the late-time relaxation, we linearize Eq. (39) around the steady state *δ* = *δ*_∞_. At the adapted steady state under a step input, we have *σ* → *σ*_*s*_ and hence *ε* → *ε*_*s*_, so the relevant tangent modulus for late-time relaxation is *𝒮*^*′*^(*ε*_*s*_). Since

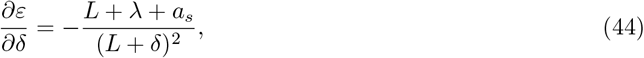

the linearized dynamics for Δ*δ*(*t*) := *δ*(*t*) *− δ*_∞_ becomes

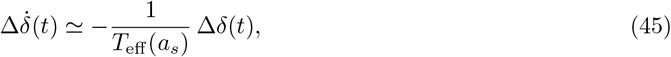

with the effective relaxation timescale

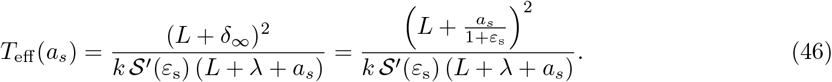

This shows that constitutive nonlinearity enters the late-time transient only through the tangent modulus *𝒮′*(*ε*_s_), while the step amplitude *a*_*s*_ contributes via the nonlinear geometry.

### Parameters used in numerical simulations

Unless otherwise stated, simulations use *L* = 5.0 *µ*m, *λ* = 5.0 nm, *E* = 2.0 GPa, *β* = 10^4^, and *k* = 0.1 nm*/*(s · kPa). The homeostatic strain and stress are *ε*_*s*_ = *λ/L* = 10^*−*3^ and 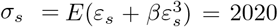 kPa. Step inputs are implemented as a smooth step with rise time *t*_rise_ = 2.0 *×* 10^*−*3^ s. The step amplitude is swept over *a*_*s*_ ∈ *{*0.06, 0.2, 0.6, 2.0, 6.0*}* nm. The simulation end time *t*_end_ is chosen automatically such that *δ*(*t*_end_)*/δ*_∞_ ≥ 0.97 for the largest *a*_*s*_. Numerical solutions use *n* = 3000 samples for the exploratory run and *n* = 5500 for the final plots.

### Numerical verification

Figure A7 shows step responses for multiple amplitudes *a*_*s*_ under the full nonlinear kinematics. The initial stress excursion increases strongly with *a*_*s*_, reflecting strain-stiffening. Figure A8 quantifies this amplitude dependence: the predicted instantaneous jump Eq. (40) agrees with the early-time numerical maximum. Because the step is smoothed for numerical stability, we compare the theoretical instantaneous jump with the early-time peak attained during the rise time. Finally, Fig. A9 shows a late-time collapse of the normalized relaxation (*δ*_∞_ *− δ*(*t*))*/δ*_∞_ when time is rescaled by *T*_eff_ (*a*_*s*_), supporting the local-linearization description. In summary, strain-stiffening amplifies the instantaneous stress excursion with step amplitude, whereas the adapted steady state and the late-time relaxation remain controlled by fixed-end kinematics and the tangent modulus evaluated at the homeostatic strain.

**Fig. A7.**
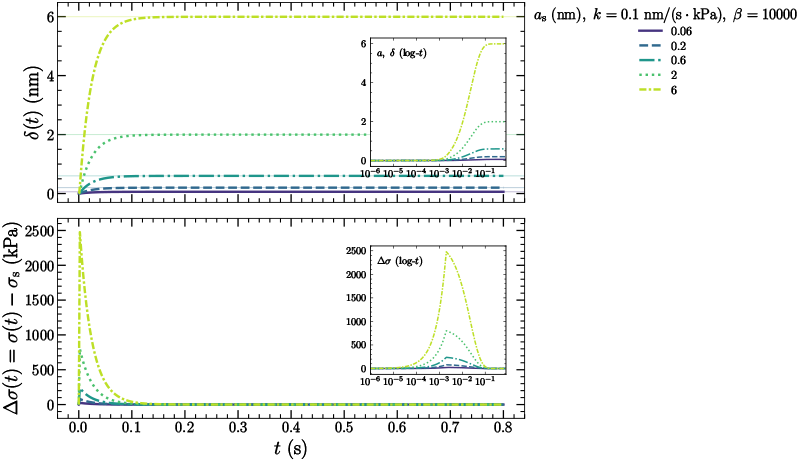
Step response of the nonlinear elastic remodeling model with strain-stiffening (full nonlinear kinematics). The imposed deformation is a step *a*(*t*) = *a*_*s*_*H*(*t*) with amplitudes *a*_*s*_ ∈ *{*0.06, 0.2, 0.6, 2, 6*}* nm (implemented as a smooth step with rise time *t*_rise_ = 2.0 × 10^*−*3^ s). The constitutive law is *σ* = 𝒮 (*ε*) = *E*(*ε* + *βε*^3^) with *β* = 10^4^ and remodeling follows 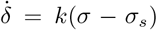 with *k* = 0.1 nm*/*(s · kPa). **(Top)** Input *a*(*t*) and rest-length change *δ*(*t*). Horizontal guides indicate the kinematic steady value *δ*_∞_ = *a*_*s*_*/*(1+*ε*_*s*_). **(Bottom)** Stress error Δ*σ*(*t*) = *σ*(*t*) − *σ*_*s*_, showing an amplitude-dependent initial stress excursion due to strain-stiffening. Insets show early-time dynamics on a logarithmic time axis.

## Funding

We thank the support from JSPS KAKENHI (24KJ1657 to EM and 23H04928 to SD).

**Fig. A8.**
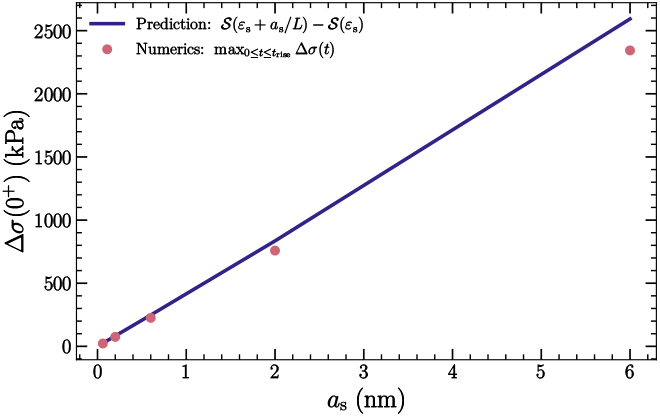
Nonlinear dependence of the step-induced stress jump on the step amplitude. The solid line shows the theoretical instantaneous jump Δ*σ*(0^+^) = 𝒮 (*ε*_*s*_ + *a*_*s*_*/L*) − 𝒮 (*ε*_*s*_). Symbols show the numerical early-time maximum 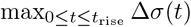 from the smooth-step implementation. The agreement confirms that the amplitude dependence of the initial stress excursion is set by the nonlinear constitutive law.

**Fig. A9.**
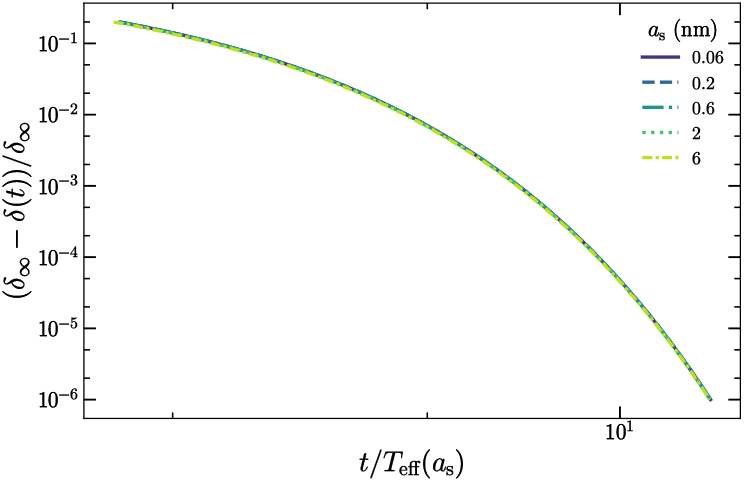
Late-time collapse of the relaxation under local linearization. Using the same simulations as Fig. A7, we plot the normalized relaxation (*δ*_∞_ *− δ*(*t*))*/δ*_∞_ against rescaled time *t/T*_eff_ (*a*_*s*_), where *T*_eff_ (*a*_*s*_) is given by Eq. (46). The collapse supports the local-linear relaxation 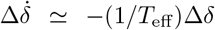 at late times, despite the amplitude-dependent initial stress jump.

## Author contributions

**Eiji Matsumoto**: Writing – review & editing, Writing – original draft, Visualization, Validation, Methodology, Investigation, Funding acquisition, Formal analysis, Data curation, Conceptualization. **Shinji Deguchi**: Writing – review & editing, Visualization, Validation, Supervision, Resources, Project administration, Methodology, Funding acquisition, Formal analysis, Data curation, Conceptualization.

